# Population genomics of two closely related anhydrobiotic midges reveals differences in adaptation to extreme desiccation

**DOI:** 10.1101/2020.08.19.255828

**Authors:** N.M. Shaykhutdinov, G.V. Klink, S.K. Garushyants, O.S. Kozlova, A.V. Cherkasov, T. Kikawada, T. Okuda, D. Pemba, R.M. Deviatiiarov, G.R. Gazizova, A.A. Penin, E.I. Shagimardanova, R. Cornette, O.A. Gusev, G.A. Bazykin

## Abstract

The sleeping chironomid *Polypedilum vanderplanki* is capable of anhydrobiosis, a striking example of adaptation to extreme desiccation. Tolerance to complete desiccation in this species is associated with the emergence of multiple paralogs of protective genes. One of the gene families highly expressed under anhydrobiosis and involved in this process are protein-L-isoaspartate (D-aspartate) O-methyltransferases (PIMTs). Recently, a closely related anhydrobiotic midge from Malawi, *P. pembai*, showing the ability to tolerate complete desiccation similar to that of *P. vanderplanki*, but experiences more frequent desiccation-rehydration cycles due to differences in ecology, was discovered. Here, we sequenced and assembled the genome of *P. pembai* and performed a population genomics analysis of several populations of *P. vanderplanki* and a population of *P. pembai.* We observe positive selection and radical changes in the genetic architecture of the *PIMT* locus between the two species, including multiple duplication events in the *P. pembai* lineage. In particular, *PIMT-4*, the most highly expressed of these *PIMTs*, is present in six copies in the *P. pembai*; these copies differ in expression profiles, suggesting possible sub- or neofunctionalization. The nucleotide diversity (π) of the genomic region carrying these new genes is decreased in *P. pembai*, but not in the orthologous region carrying the ancestral gene in *P. vanderplanki*, providing evidence for a selective sweep associated with post-duplication adaptation in the former. Overall, our results suggest an extensive recent and likely ongoing, adaptation of the mechanisms of anhydrobiosis.

## Introduction

The anhydrobiotic midge *Polypedilum vanderplanki* (Chironomidae) (fig. S1a) in natural habitat shows a unique example of adaptation to desiccation. The larval stage of the midge inhabits temporary pools formed during rains on granite boulders in the northern part of Nigeria. During the dry season, the larvae lose 99.2% of water, replacing it with trehalose combined with other molecular protectants, that allows the larva to survive the dry period in the dried state, without detectable metabolism. After rehydration with the start of the rainy season, the larva returns to active life within less than an hour (Sogame and Kikawada 2017). Remarkably, the chironomids generally demonstrate high ecological plasticity, and their larvae are known for their ability to adapt to a wide range of extreme conditions, including high salinity, anaerobic environment, low pH, low or high temperatures or desiccation (Armitage et al. 2012). For instance, another extremophilic midge *Belgica antarctica* that lives in the Antarctic can survive water loss of up to 70%, sustaining the low temperatures of the Antarctic climate (Kelley et al. 2014).

Physiological aspects of anhydrobiosis in the sleeping chironomid were investigated since the early 1950s (Hinton 1951), but studies of mechanisms underlying anhydrobiosis were launched only after the establishing of a successful rearing protocol for this species (Watanabe et al. 2002). Recent work on the molecular mechanisms underlying anhydrobiosis has led to the identification of several groups of biological molecules that contribute to resistance to desiccation in *P. vanderplanki*. These include the protein-L-isoaspartate (D-aspartate) O-methyltransferases (PIMTs), proteins involved in trehalose biosynthesis, late embryogenesis proteins (LEA proteins), antioxidant system proteins, heat shock proteins (HSP) and DNA repair enzymes (Kikawada et al. 2006; Cornette and Kikawada 2011; Gusev et al. 2011). Comparison of *P. vanderplanki* to a non-desiccation-tolerant relative *Polypedilum nubifer* showed that genes encoding desiccation-specific proteins (*PvLEA*, *PvTRX*, *PvPIMT*, *PvHb*) in *P. vanderplanki* were present in multiple copies, highly transcribed and located in compact genomic clusters called Anhydrobiosis-Related gene Islands (ARIds) (Gusev et al. 2014).

One of the most interesting gene families associated with anhydrobiosis is *PIMT* genes, which have multiplied to 14 copies in the *P. vanderplanki* genome. This group of genes is remarkable in that they have one of the strongest changes in expression level in response to desiccation of the midge larva among all genes (Gusev et al. 2014); however, their molecular role in anhydrobiosis is not well understood (Deviatiiarov et al. 2017). The enzyme encoded by *PIMT* belongs to the group of S-adenosylmethionine (SAM) dependent methyltransferases and catalyzes the repair of damaged amino acids such as L-isoaspartate and D-aspartate (Khare et al. 2011). Genes encoding these enzymes are highly conserved and are present as a single copy in genomes of all eukaryotes (including insects), archaea and gram-negative eubacteria, to the exception of plants and some bacteria that have several *PIMT* isoforms (Desrosiers and Fanélus 2011). The activity of PIMT in animals is associated with resistance to stress factors and is directly related to life expectancy (Desrosiers and Fanélus 2011; Khare et al. 2011). It was shown that the accumulation of PIMT1 protein in *Arabidopsis thaliana* was associated with reduced accumulation of abnormal L-isoaspartyl residues and thus lead to increased longevity and vigor of dried seeds (Ogé et al. 2008).

A family of *P. vanderplanki* genes, *PvPimt*, are classified as *PIMTs* based on the presence of the conserved PIMT functional domain in the proteins encoded by them. Desiccation of larvae is associated with a significant increase in transcription of genes of this family (Gusev et al. 2014), indicating that it plays an important functional role in anhydrobiosis. However, individual *PvPIMTs* demonstrate marked differences in their amino- and carboxyterminal regions, suggesting that they may differ in their preferences for substrates, localization, or other specific properties (Gusev et al. 2014).

While other lineages (tardigrades, rotifers, nematodes) have many anhydrobiotic organisms in their phyla, *P. vanderplanki* was untill recently the only proven anhydrobiotic species both among the Chironomidae family and the whole phylum Arthropoda (Watanabe 2006). However, another midge from Malawi that was initially referred to as *P. vanderplanki* (McLachlan 1983) is also able to survive in the desiccated state in the larval stage but differs in ecology from the Nigerian anhydrobiotic midge (Cornette et al. 2017). While in Nigeria the dry season can last up to eight months without rain, in Malawi, it is interspersed with periodic sporadic rainfall (Cornette et al. 2017). Therefore, the larvae of the Malawian midge face several desiccation-rehydration cycles within one season (Cornette et al. 2017). Together with morphological differences between the two chironomids, this allowed to describe the Malawian midge as a new anhydrobiotic species *Polypedilum pembai* (Cornette et al. 2017) (fig. S1b). Comparative analysis of genomic data from the two described anhydrobiotic species belonging to the same genus can help identify adaptations to desiccation at genome and transcriptome levels and understand how these species evolved ability for anhydrobiosis.

In this study, we compare genomes of populations of *P. vanderplanki* and *P. pembai* to study the evolutionary adaptations to anhydrobiosis. We detect past duplication events leading to an increase in the numbers of desiccation response paralogs *(PIMTs)* in the lineage of *P. pembai (PpPIMTs).* Specifically, *PIMT4*, the paralog which is the most transcribed in response to desiccation among all *PIMTs* genes in both midges, is present in multiple copies in the genome of *P. pembai.* We find that the copies of *PpPIMT4* have experienced positive selection, suggesting that adaptation to anhydrobiosis is ongoing.

## Results

### Assembly and characteristics of the anhydrobiotic midges genomes

We obtained and sequenced seven populations of midges: six populations of *P. vanderplanki* from six sampling sites in Nigeria (fig. 1b) and one population of *P. pembai* from one sampling site in Malawi (see Materials and Methods). The summary statistics for the samples are shown in table S1.

**Fig. 1.**
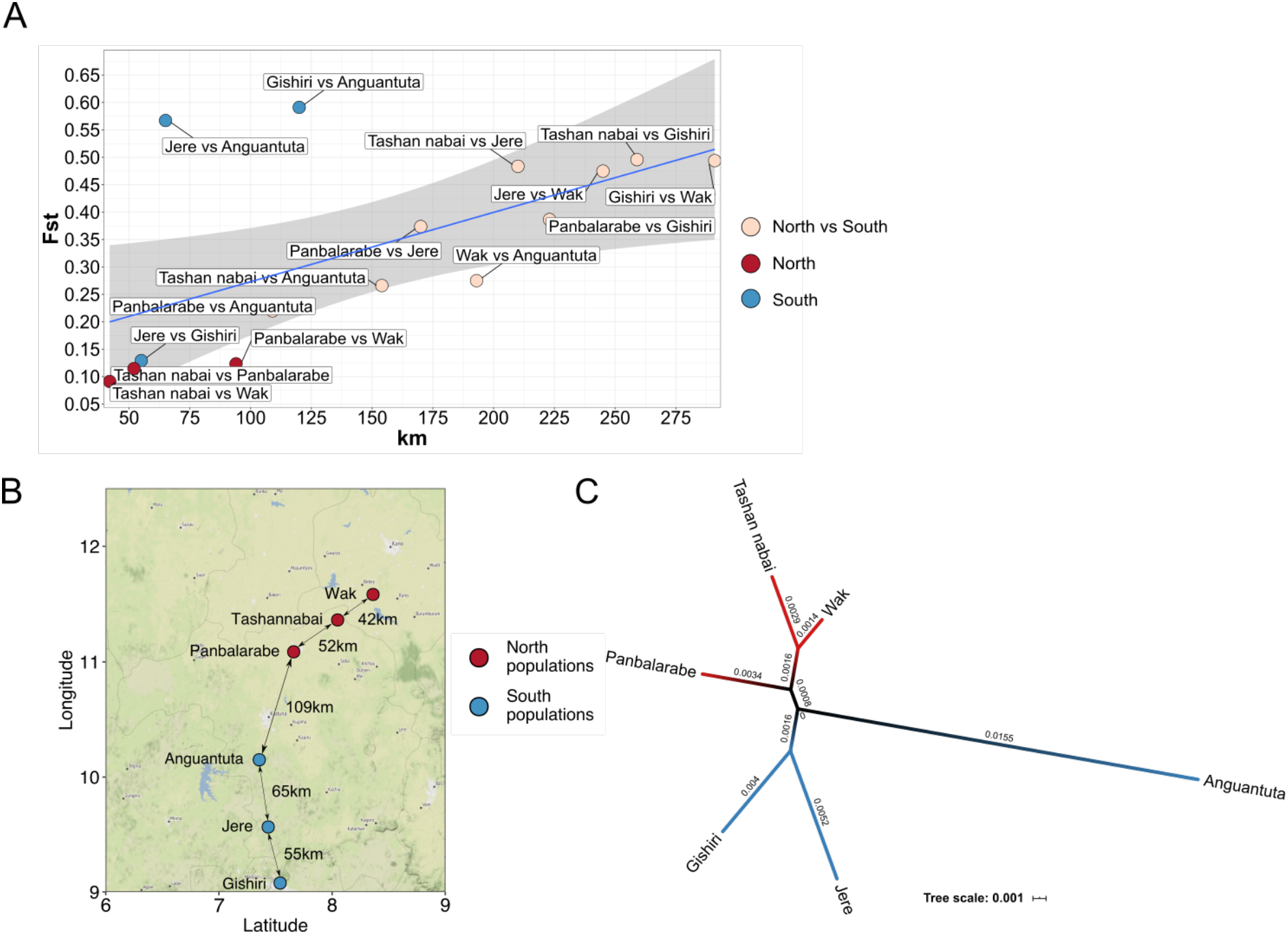
Geographic structure of *P. vanderplanki* populations. (A) Genetic distance as a function of geographic distance. (B) Map of collection sites of *P. vanderplanki* populations. (C) Unrooted ML tree based on concatenated mtDNA protein-coding genes. Blue color represents the northern cluster of populations; red color represents the southern cluster of populations. Values on branches represent branch lengths.

As a reference genome we used a reassembly of the *P. vanderplanki* genome, details of which will be published separately (Yoshida et al., in preparation). In brief, this assembly represents a complete genome with four chromosomes (N50 = 35 Mb, which is the length of the second chromosome) and 384 non-chromosomal scaffolds. The size of the genome assembly is about 119 Mb, and the GC content is 28%. According to BUSCO, the assembly completeness of the *P. vanderplanki* genome relative to the *Diptera* Odb10 dataset (3285 proteins) is 95.7% (including 94.7% as single-copy proteins), with 3.6% of the reference proteins missing. Based on a k-mer analysis, the average coverage of *P. vanderplanki* genome by Illumina-derived reads was estimated as 375x, coverage by long reads was about 10x (Kozlova et al., unpublished data).

The draft genome of *P. pembai* was assembled de novo from *P. pembai* sample. About 57 mln shotgun paired-end reads (100 bp) were used for draft genome assembly of *P. pembai*. The genome size of this species was estimated at 0.096-0.109 pg DNA as 94-107 Mb, which is roughly the same size as *P. vanderplanki* (Cornette et al. 2015). The estimated average coverage of the *P. pembai* draft genome is between 95x and 114x, and the final size of the draft genome assembly is ~122Mb. According to BUSCO, assembly completeness of *P. pembai* genome (by the same *Diptera* dataset) was estimated as 94.8% (92.7% for single-copy proteins), and 4.2% of the reference proteins were missing. The basic structural and functional features of *P. vanderplanki* and *P. pembai* genome assemblies are presented in table S2.

Additionally, we obtained high-throughput total RNA-seq data for *P. pembai* from midge larvae in the wet state and after desiccation for 24h and 48h. Total RNA-seq data of *P. pembai* was used to compare transcription responses to desiccation of *P. pembai* and *P. vanderplanki* larvae. Each sample was prepared in two technical replicates. A high rate of unmapped reads can be explained by a large number of rRNA reads (table S3).

### Population genomics of *P. vanderplanki*

We collected individuals of the adult stage of *P. vanderplanki* from six sampling sites in Nigeria (Materials and methods) (fig. 1b). Sampling sites were tens to hundreds of kilometers from each other, which could limit gene flow between midge populations. Within-population gene diversity (π) varied between 0.004 and 0.007 for different populations of *P. vanderplanki* (table S4). Pairwise genetic distances between populations were higher: between 0.005 and 0.008 in the comparisons of northern populations, between 0.005 and 0.015 in the comparisons of southern populations, and between 0.009 and 0.014 in the north-south comparisons (table S4, fig. S2).

To better understand the genetic structure of the six considered populations, we analyzed the Wright fixation index (*Fst*). *Fst* was above-zero in all comparisons, varying between 0.114 and 0.495 (fig. 1a), indicating deviation from panmixis. It was lower, indicating weaker isolation, in the south-south and north-north comparisons (between 0.09 and 0.13, to the exception of Jere vs. Anguantuta and Gishiri vs. Anguantuta comparisons, for which it equaled 0.56 and 0.59 respectively) than in the south-north comparisons (between 0.22 and 0.49). Consistently, *Fst* was correlated with geographic distance between the analyzed populations (Mantel test R = 0.575; p-value = 0.0375) (fig. 1a), especially when the outlying Anguantuta population was excluded from analysis (Mantel test R = 0.954; p-value = 0.025).

To confirm geographic subdivision of *P. vanderplanki* populations, we estimated phylogenetic distances based on concatenated sets of mitochondrial protein-coding genes (fig. 1c). Such data provide a higher phylogenetic resolution, compared to the frequent approach of using just a single mtDNA gene such as COI (cytochrome c oxidase I) which often does not allow genetic differentiation of closely located populations (Havird and Santos 2014). In line with the *Fst* analysis, we observe that the six populations form two distinct clusters - the northern and the southern and that the Anguantuta population is an outlier, positioned at a high evolutionary distance from the remaining northern populations (fig. 1c). Morphological analysis did not show significant differences between the populations, although the superior volsella of male genitalia, which is an important taxonomic feature, showing a lateral seta in 10-20% of the individuals from Wak, Tashan nabai and Panbalarabe, whereas such seta was never observed in southern populations (supplementary table S5). A slightly higher male antennal rate in the southern populations seemed also to support the genetic separation between northern and southern populations.

Based on the observed geographic subdivision of *P. vanderplanki* populations, we hypothesized that the mechanisms of anhydrobiosis encoded by the *PIMT* gene family could have evolved to adapt to their specific microenvironments. However, we found no support for this hypothesis either at the level of copy number variation or single-nucleotide variation of anhydrobiosis-related genes. Indeed, the coverage of individual *PIMT* paralogs involved in anhydrobiosis was similar between populations in our Pool-seq data, indicating that they were present in the same number of copies in different populations (supplementary table S6). Similarly, the estimated πN/πS ratio of *PvPIMT* genes (supplementary table S7) was low (πN/πS < 0.5) across all six *P. vanderplanki* populations, indicating the action of negative selection on all gene copies; we saw no evidence of relaxed negative selection or positive selection. Together, these findings indicate that the aspects of anhydrobiosis associated with *PvPIMT* genes are conserved within and between the populations of *P. vanderplanki*.

### Comparative genomics of *P. vanderplanki* and *P. pembai*

While *P. vanderplanki* was the only anhydrobiotic insect known until recently, the discovery of another closely related species, *P. pembai*, changed this view (Cornette et al. 2017). *P. pembai* and *P. vanderplanki* are closely related according to the COI gene data (Cornette et al. 2017); however, morphological and cytological data (Cornette et al. 2017) indicate that they are two distinct species. Reproductive isolation is often associated with genetic distance above ~10% (Mendelson et al. 2004; Elliot and Crespi 2006). Our analysis showed strong genetic differentiation between *P. vanderplanki* and *P. pembai* (*Fst* = 0.85 to 0.91, genetic distance varied between 0.077 and 0.08 depending on the considered *P. vanderplanki* population; table S8), suggesting reproductive isolation between these species and supporting their species status. Within-population gene diversity (π) for *P. pembai* equaled 0.004, similar to within-population gene diversity (π) for *P. vanderplanki* populations.

Given the closeness of *P. vanderplanki* and *P. pembai*, it is unlikely that they developed anhydrobiosis independently. Consistently with this assumption, we observe a similar genetic architecture of anhydrobiosis-associated genes, in particular, the presence of a multigene family of *PIMTs* (see below). Still, due to the difference in the ecotopes of *P. pembai* and *P. vanderplanki*, the frequency of desiccation-rehydration cycles is expected to be higher for *P. pembai (Cornette et al. 2017)*, suggesting the possibility of a species-specific adaptation. Indeed, the genetic distance between *P. vanderplanki* and *P. pembai* was higher for chromosome 4 (10%) than for the other three chromosomes (7%) (table S8). Chromosome 4 carries the majority of anhydrobiosis-related genes, including *PvPIMT2-14*, in *P. vanderplanki*, so increased divergence may indicate accelerated evolution of genomic features connected with the ability to survive desiccation. We asked whether these differences have led to any observable differences in the genetics of anhydrobiosis between the two species.

### Adaptive evolution of the *PIMT* gene family in *P. pembai*

Comparative genomic analysis revealed the presence of 14 paralogs of *PIMT* in *P. vanderplanki*, and 19 paralogs in *P. pembai.* Using transcriptome data, we found that all 19 paralogs are differentially expressed in *P. pembai* in response to desiccation. The transcriptome profile of desiccation response in *PvPIMTs* and *PpPIMTs* is very similar between the two species. In particular, the two paralogs most highly expressed in response to desiccation, *PpPIMT4-1* (~12500 RPKM at D48) and *PpPIMT12*, are the two most highly expressed paralogs as *PvPIMT4* (>28000 RPKM at D48) and *PvPIMT12*) (fig. 2). The expression rate of the other copies of *PIMT4* in *P. pembai, PpPIMT4-2* to *PpPIMT4-6*, remained low in response to desiccation but was still elevated till complete desiccation (D48). Overall, the transcriptional profile of *PIMT* was conserved between the two species (fig. 2), despite the fact that the two species have diverged ~ 49 million years ago (Cornette et al. 2017).

**Fig. 2.**
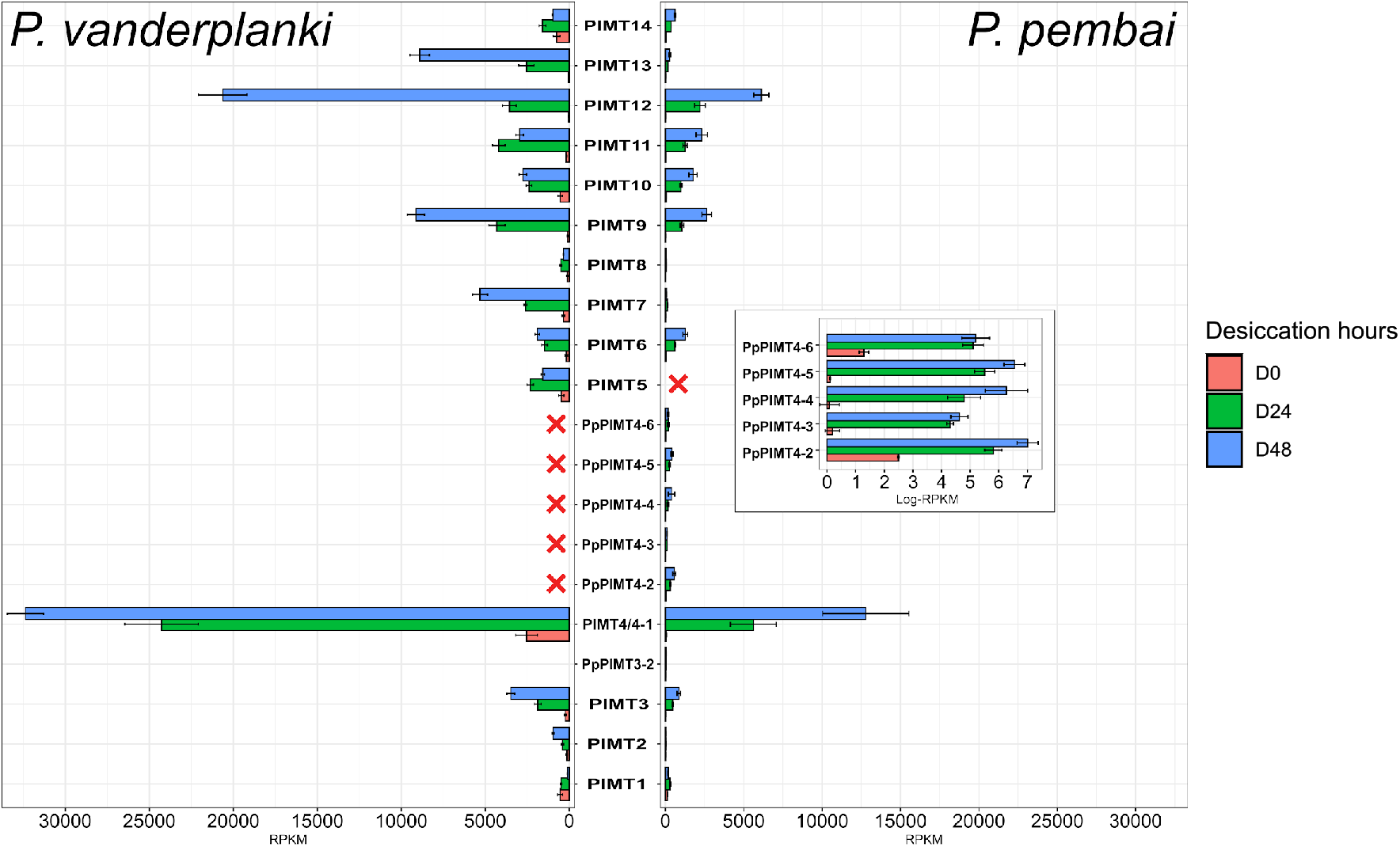
Differential expression of *PIMTs* in *P. vanderplanki* and *P. pembai* in response to desiccation. The expression profile of 14 *PvPIMTs* (left panel) and 19 *PpPIMTs* (right panel) during desiccation. The inset in the right panel represents the expression of duplicated paralogs in log scale. Red crosses identify paralogs absent in the corresponding genome. D0: control, D24, D48: desiccation for 24 and 48 h, respectively. Pv, *P. vanderplanki;* Pp, *P. pembai.*

Given the overall high conservation of the PIMT locus, we were interested in the difference in the number of *PIMT* copies between the two species. Phylogenetic analysis of the *PIMT* family indicates that this difference is due to a prolific clade of *P. pembai* paralogs in the PIMT gene family tree: one of the *P. vanderplanki* genes, *PIMT4*, is present in six copies, *PIMT4-1* to *PIMT4-6*, in *P. pembai* (fig. 3a). Analysis of genomic positions of *PIMTs* in the two species indicates that the order of genes is generally well-preserved between them; the increase in the size of the locus from 60 kb in *P. vanderplanki* to 85 kb in *P. pembai* is due to the presence of additional copies in *P. pembai* (fig. 3c). *PIMT4-1* to *PIMT4-6* are positioned in tandem in *P. pembai*, and their positions are syntenic to that of *PIMT4* in *P. vanderplanki* (fig. 3c), supporting their orthology.

**Fig. 3.**
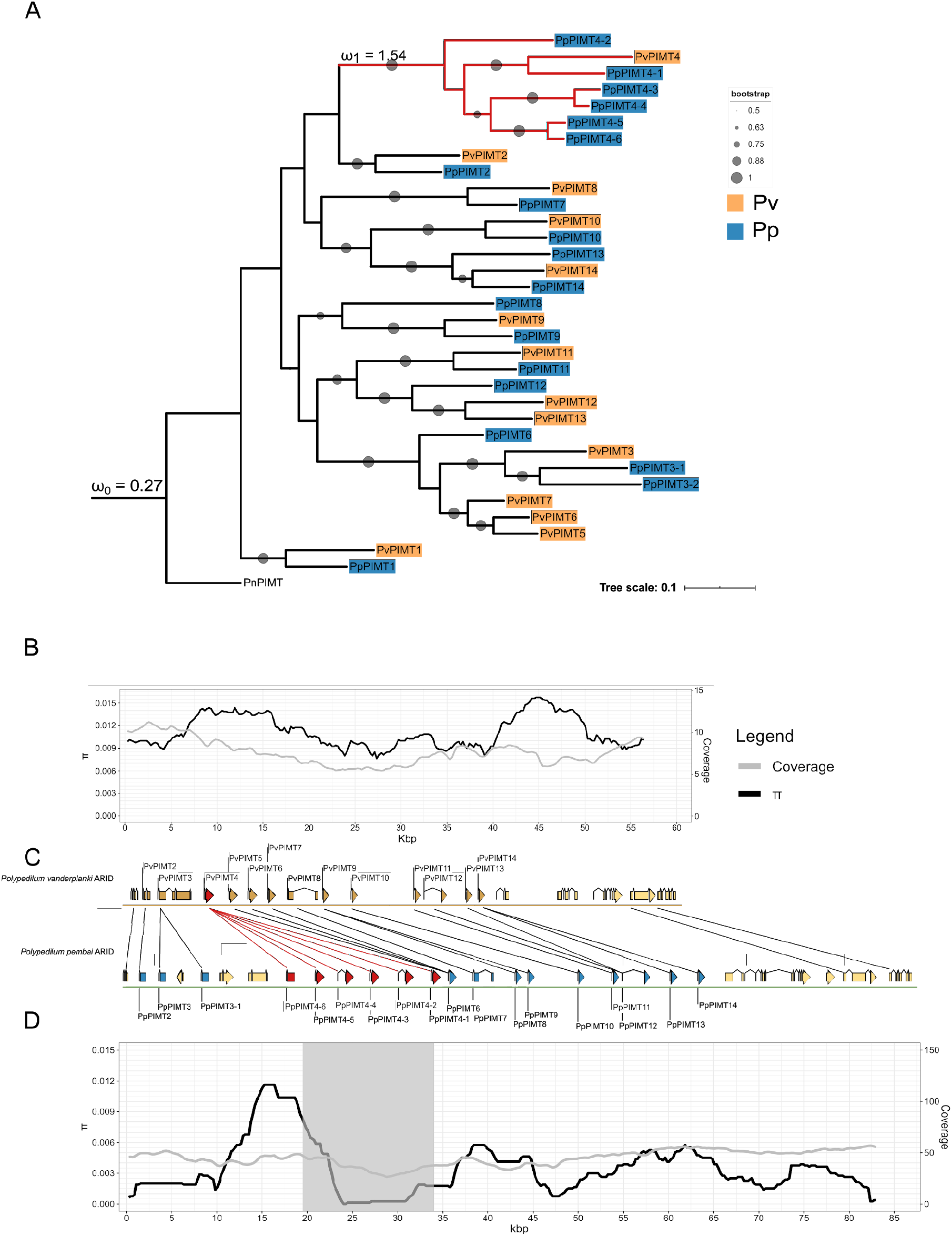
Comparative analysis of the *PIMT* ARID from two midges. (A) ML tree with bootstrap testing (100 replicates). Pv, *P. vanderplanki*; Pp, *P. pembai*, Pn, *P. nubifer*. Red color shows the clade under positive selection; blue color shows *P. pembai PIMTs*; orange color shows *P. vanderplanki PIMTs.* (B) Nucleotide diversity along *P. vanderplanki PIMT* ARID; (C) Comparison of gene order in the two ARID. Black lines represent homologous genes; (D) Nucleotide diversity along the *P. pembai PIMT* ARID.

The phylogenetic branches leading to four of these genes, *PIMT4-3* to *PIMT4-6*, are shorter than those corresponding to the *P. vanderplanki - P. pembai* divergence (fig. 3a), indicating that at least some of these differences in copy number between species were caused by a duplication in the *P. pembai* lineage rather than a gene loss in the *P. vanderplanki* lineage. To formally ask when this divergence has occurred, we calibrated the phylogenetic tree using fossils data (Cornette et al. 2017), and used RelTime-ML model (Tamura et al. 2012) to date the duplication events. This analysis suggests that the amplification of this gene family has occurred in 5 duplication events, dating to 53.28, 51.82, 39.71, 8.89 and 7.26 mya (fig. S3). The earliest two of these events predate the *P. vanderplanki - P. pembai* divergence, indicating that subsequent evolution could have involved a gene loss in *P. vanderplanki;* conversely, the three later events have unambiguously occurred in the *P. pembai* lineage.

We hypothesized that this amplification of the *PIMT* gene family has triggered adaptive evolution. Consistent with this hypothesis, we observe a reduction in nucleotide diversity in the genomic region containing the additional paralogs of *PIMT4* in *P. pembai* (fig. 3d). This difference in diversity from neighboring regions is picked up by Pool-hmm (Boitard et al. 2013) as evidence for a past selective sweep in this region. No such reduction is observed in the homologous region of *P. vanderplanki* (fig. 3b), indicating that positive selection has only affected one of the two diverging species.

Given the observed trace of a selective sweep, we asked whether the *PIMT* gene family demonstrates evidence for positive selection at the codon level. To address this, we first measured the pairwise dn/ds values between all pairs of *P. vanderplanki* and *P. pembai PIMT* genes (i.e., *PvPIMT1* vs. *PpPIMT1*, *PvPIMT1* vs. *PpPIMT2*, etc.). The observed distribution of dn/ds values had a mean of 0.33 (fig. 4a). To get a reference point for this value, we observed that the mean dn/ds value depends on the number of genes in the orthogroup, with smaller orthogroups having lower dn/ds. Therefore, for comparison, we used the set of orthogroups in *P. vanderplanki* - *P. pembai* comparisons that had >7 genes in each species; there were 69 such orthogroups (supplementary fig. S4). The observed dn/ds values for the *PIMT* gene family was higher than that for other orthogroups with >7 genes (0.33 vs. 0.30, p-value < 0.0007, Mann-Whitney U Test; fig. 4a). This indicates that the *PIMT* gene family evolves somewhat more rapidly than expected for an orthogroup of its size, suggesting relaxation of negative and/or presence of positive selection.

**Fig. 4.**
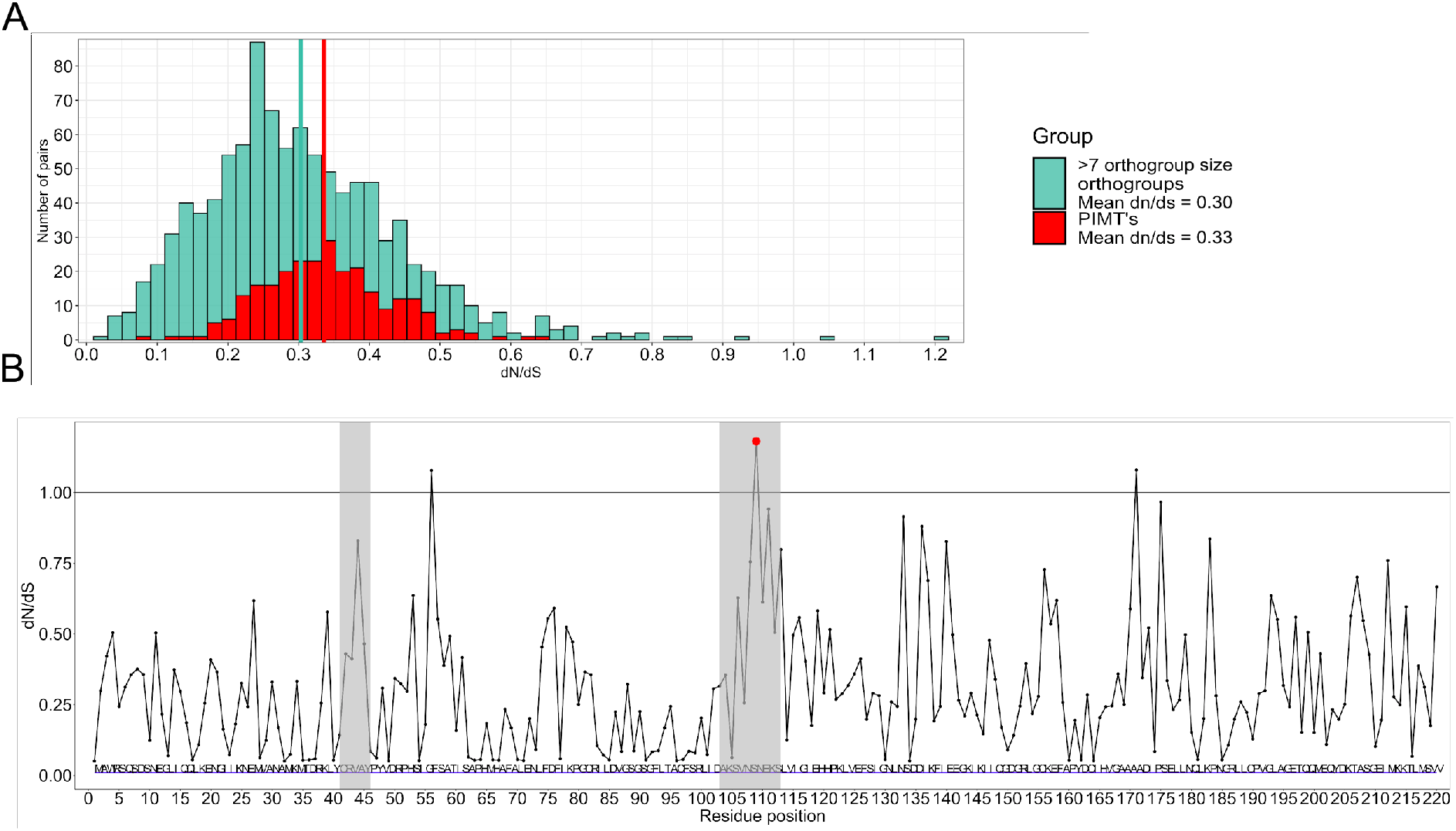
Selection at *PIMT* genes. (A) Distribution of dn/ds values of >7-sized orthologous groups and PIMTs genes. The turquoise line represents the mean of the dn/ds distribution of >7-sized orthologous groups; the red line represents the mean of the dn/ds distribution of *PIMTs* genes. (B) Identification of sites in *PIMTs* genes under positive selection. The only residue with BEB score >50 is shown in red. Grey boxes indicate the two short regions specific to the *PIMTs* of *Polypedilum sp.*

To better understand the patterns of selection in *PIMT* genes, we asked how selection differs between protein sites. For this, we applied the site test for positive selection to the codon alignment of the *PIMT* gene family, using the M2a (selection) and M8 (beta & ω) models implemented in codeml. We found several sites with a statistically significant signal of positive selection (BEB score higher than 50%; fig. 4b). The sites with ω > 1 are positioned in two regions of the *PIMT* alignment: sites 41 to 46 and 103 to 113. These sites do not overlap the *D. melanogaster* model of its only methyltransferase (Deviatiiarov et al., in preparation).

Finally, given our observation of a selective sweep associated with the increase in the *PIMTs* copy number, we hypothesized that the subsequent evolution of the duplicated region could also be adaptive, in line with the neofunctionalization models (Innan and Kondrashov 2010). To test this, we used the branch test of codeml to compare the ω values in the clade that has expanded in the *P. pembai* lineage (foreground model, ω1) to those in the rest of the *PIMT* tree (background model, ω2). Strikingly, we observe a strong gene-wide signal of positive selection in the duplicated clade (ω1 = 1.5), while the rest of the tree evolves under strong negative selection (ω0 = 0.3; p-value < 0.001; fig. 3a). This indicates that the post-duplication amino acid-changing mutations in the PIMT genes conferred a selective advantage.

## Discussion

The anhydrobiotic midges represent a unique example of adaptation to extreme conditions. Here, we provide the first comparative genomics analysis of two species of anhydrobiotic midges. By sequencing multiple populations of *P. vanderplanki*, we detect a high degree of genetic structure associated with geographic distance, indicating population subdivision. Such a geographic differentiation is perhaps unexpected for a flying insect, and appears to be stronger than in other studied Diptera (Kumar and Singh 2017), despite the lack of geographic or ecological barriers between our six populations. However, limited flight in *Polypedilum sp.* is consistent with other data. The maximal dispersal distance evaluated for *P. pembai* (formerly identified as *P. vanderplanki*) in Zomba, Malawi, ranged between 0 and 446 m, suggesting a poor flight ability for this insect (McLachlan 1983). In addition, many chironomid species stop swarming under windy conditions in order to avoid being blown away from their habitat (Armitage, 1995). Thus, dispersal of the anhydrobiotic midges with the winds on larger distances should be accidental and this corroborates isolation of the populations, explaining the high Fst values observed here between Nigerian locations distant from more than 50 km.

Geographic isolation creates ample opportunity for lineage-specific adaptation. According to fossil data, the lineages of *P. vanderplanki* and *P. pembai* diverged 48 mya. Our analysis indicates that these lineages have accumulated ~8% genetic distance, corresponding to the substitution rate of ~10^−9^ per nucleotide per year which is consistent with other *Dipterans* (Keightley et al. 2014). While these species are both capable of anhydrobiosis, differences in their ecology could lead to differences in evolutionary pathways of this system.

Most notably, we describe the amplification of the gene family encoding an important protein involved in anhydrobiosis, and provide evidence that the genes of this family have accumulated adaptive amino acid substitutions after duplication. Specifically, we reveal the presence of positive selection acting on additional copies of methyltransferases in *P. pembai* (fig. 3a), as well as the presence of a relaxed selection acting on *PIMT* genes involved in anhydrobiosis (fig. 4a).

Gene duplication is one of the main sources of functional gene diversity. It is believed that one of the main factors in the formation of paralogs in the genome is adaptation to changing environmental conditions (Kondrashov 2012). In the Antarctic notothenioid fish (*Dissostichus mawsoni*) multiple duplications of the gene family of antifreeze glycoprotein (AFGP) occurred, based on the paralogue of trypsinogen in response to the cold habitat of the fish (Chen et al. 1997). It is noteworthy that duplication of the original trypsinogen and the appearance of the first antifreeze gene occurred 5-14 million years ago, which roughly coincides with the time of mid-Miocene (10–14 mya), when the Antarctic Ocean started to freeze (Chen et al. 1997).

Here, we provide evidence that the duplication of *PIMT* genes in *P. vanderplanki* and *P. pembai* genomes has been adaptive, possibly allowing the evolving *P. pembai* lineage to adapt to the changing environment. An increase in the copy number of *PIMT* from 1 to 14 copies suggests neofunctionalization of these genes, which is confirmed by large distances between the paralogs, and most importantly, the absence of methyltransferase function in them (Deviatiiarov et al., in preparation). *PIMT1* and *PIMT2* from both species have low expression during anhydrobiosis, unlike other paralogs that have increased expression (fold change D48 vs D0 > 12 for *PvPIMT4*) in response to desiccation. Despite similarity with the model of gene amplification - beneficial increases in dosage (Innan and Kondrashov 2010), the functions of *PIMTs* paralogs differ from the functions of the founder ancestor.

At the moment, the biological role of *PIMT* paralogs in cell during desiccation is not known exactly; therefore, the significance of their role can be assumed from the result of evolutionary analysis and from transcriptome profiles (fig. 3–4) at various desiccation time points. While the gene-wide dn/ds ratios remained below 1, indicating ongoing negative selection, the relaxation of this selection in the two ARIDs suggests the possibility of evolution towards anhydrobiosis. Also, the fact that the branch model analysis of dn/ds has detected a positive selection on the branch with copies of PpPIMT4 suggests that *P. pembai* midge still continues to adapt to extreme living conditions.

We don’t know if duplication itself has been adaptive - just the subsequent evolution. However, the high rate of amplification indicates that this very likely has been the case. The *PIMT* gene has remained conserved and single-copy over the course of billions of years, but has rapidly undergone five duplication events in just one lineage over the period of ~ 50 million years - a fact hardly consistent with the neutrality of these duplication events.

Duplications of *PIMT4* paralog in *P. pembai* can be described by a model of gene duplication with a modified function when positive selection acts on duplicates (Innan and Kondrashov 2010). The following observations support in favor of chosen model: the presence of a positive selection acting on these paralogs (fig. 3a) and selective sweep in the *PIMT4* paralog region (22.5kbp - 35kbp) containing 4 of 6 copies of *PIMT4*, which can indicate the rapid evolution of *PpPIMT* copies. The only contradiction in the choice of the model is that *PpPIMT4* is strongly expressed in response to desiccation (fig. 4), whereas the paralogs from *PpPIMT4-2* to *PpPIMT4-6* do not have strong expression. The increased expression of the founder ancestor and the fact that PIMT genes belong to stress genes indicates a similarity with the beneficial increases in dosage model. This contradiction between the selection of models can only be resolved by experimental confirmation of the new functions of the *PpPIMT4* paralogs.

Also, the fact that these copies are under positive selection rejects the assumption that they are just duplication polymorphism. Using *PIMT4* as an example, we can assume that the founding ancestor of paralogs could have an increased tendency to duplicate if its functional significance is great. By the functional significance of a gene, we mean increased expression. Thus, we can assume that one of the signs that a gene can be duplicated is increased expression of this gene under stressful conditions, which for our model organism is desiccation stress.

However, what factor leads to the fixation of the paralogs is debatable. The divergence level between paralogs can be affected by processes such as gene conversion or adaptive introgression, so our estimates of the timing of duplication events, which are based on divergence levels, should be taken with caution. Nevertheless, divergence levels between paralogs imply that the amplification of the PIMT family has not occurred as a single event, but instead has spanned a long period of time between ~60mya and ~5mya. What happened in Africa over this time frame?

We suggest that the time of occurrence of paralogs in the anhydrobiotic midges should coincide with the time of climate change towards aridity. After 33 MYA, the Oligocene was marked by general aridification in Africa, with, in particular, desertification of South Africa, and then during the early Miocene tropical rainforests were reduced, leaving a large proportion of the African continent covered with grassland and semi-arid savannah (Feakins and Demenocal 2010). Actually, the time range of divergence between *P. vanderplanki* and *P. pembai* is overlapping with the Oligocene epoch (Cornette et al., 2017). Most likely, the origin of additional copies of methyltransferases can be explained by recent differences in the climate between a less dry Malawi and a more arid Nigeria. A more frequent occurrence of rain during the dry season in Malawi leads to the fact that *P. pembai* larvae can experience several desiccation-rehydration cycles, in contrast to generally one desiccation-rehydration cycle per dry season for *P. vanderplanki* larvae. Such a stress factor can both increase the limiting selection and lead to consolidate new beneficial mutations, including paralogs.

In summary, we show that two anhydrobiotic midges adapted to anhydrobiosis similarly the same way, they have the same gene families that are organized in clusters of paralogs. These anhydrobiosis related genes are differentially expressed as in *P. vanderplanki*. We can say that additional copies of PIMTs that we found in *P. pembai* genome can be an example of ongoing or recent events of adaptation to anhydrobiosis. Unfortunately, we cannot say anything about the molecular functions of these additional copies, but presence of expression in response to desiccation process can tell us about their potential importance in adaptation to anhydrobiosis.

## Materials and Methods

### Material collection and genome sequencing

In this work were used samples of *P. vanderplanki* collected into temporary rock pools in the semi-arid territories of the northern region of the Federal Republic of Nigeria. The sampling sites correspond to six locations (Tashan nabai, Wak, Panbalarabe, Jere, Gishiri, Anguantuta) and are plotted on a map (fig. 1b). Names of samples (populations) match the names of villages where they were sampled. Samples of *P. pembai* were collected in the central and eastern regions of the Republic of Malawi. Larvae were obtained from seasonal rock pools in Zomba city, in a location called Chikopa (15^∘^ 23^′^ 422S, 35^∘^ 18^′^ 877E) as described previously (Cornette et al. 2017). Desiccated larvae were brought back to Japan and stored in a desiccator until rehydration prior to use. Rehydrated larvae were reared on milk-agar diet as described earlier (Watanabe et al. 2002) and developed to pupae and adults, which were collected for the extraction of nucleic acids.

Transcriptomic data of *P. vanderplanki* was obtained from genome browser ‘MidgeBase’ (http://bertone.nises-f.affrc.go.jp/midgebase) and consisted of three duplicated experimental points (D0, D24, D48).

Genomic DNA was extracted from 8 to 12 adult individuals (imago) from every population. Imago were homogenized with polypropylene pestle in Eppendorf plastic tubes (1.5 mL). Highly pure genomic DNA extraction was performed with NucleoSpin Tissue kit (Clontech Takara), according to the instructions of the manufacturer. Genomic DNA concentration was estimated using a Qubit™ 3.0 fluorometer (Invitrogen™) with Quantifluor dsDNA system (Promega).

Thus, each DNA pool was a collection of samples from a specific midge population. Then, gDNA was fragmented using Covaris s220 (USA) DNA shearing protocol. The length of DNA fragments was estimated using Agilent Bioanalyzer 2100 (Agilent technologies). Libraries from each pool of gDNA were prepared using NEBNext® Ultra™ II DNA Library Prep Kit for Illumina® following manufacturer’s protocol. The concentration of libraries was measured by Qubit™ 3.0 fluorometer (Invitrogen™), its quality was verified on the Bioanalyzer using DNA High Sensitivity chip (Agilent technologies). Before sequencing, the number of molecules in each library was validated by real-time PCR using 2.5x Reaction mixture for PCR-RV in the presence of EVA Green (SINTOL, Russia) and primers for Illumina adapters (Evrogen, Russia). Further, taking into account the actual molar concentration of each library, they were diluted to 2 nM and pooled in accordance to the sequencing depth. Final pool was diluted to 11 pM.

Sequencing was carried out on the HiSeq 2500 platform (Illumina) in the pair-end mode using HiSeq Rapid Pair-end Cluster Kit v2 and HiSeq Rapid SBS Kit v2 500 cycle kit (Illumina) reagents. Agilent 2100 Bioanalyzer (Agilent Technologies, Santa Clara, CA) was used to obtain library lengths. Qubit 2.0 (Life Technologies) and real-time PCR were used to quantify libraries. Diluted libraries (with final concentration 9pM) were clustered using cBot instrument (Illumina) with TruSeq PE Cluster Kit v3 (Illumina) and after that sequenced using Illumina HiSeq 2000 sequencing machine. Length of forward and reverse reads were 250 bp and 100 bp for Anguantuta population in libraries made from Nigerian midge gDNA. Mean length of insertion varied from 227 bp (for Anguantuta population) to 418 bp (for Gishiri population). Length of forward and reverse reads were 100 bp in WGS libraries from *P. pembai* midge population gDNA. Mean length of insertion varied from 200 bp to 256 bp from *P. pembai* populations.

Concerning transcriptomic data, total RNA was obtained from larvae of the genomic strain of *P. vanderplanki*, which is an inbred line obtained from an original mixture of some populations cited above. RNA-seq data were obtained from wet larvae and larvae desiccated for 24h and 48h, as described previously (Gusev et al., 2014). Concerning *P. pembai*, larvae from Chikopa population in the wet state or desiccated for 24h or 48h as described previously (Watanabe et al., 2002) were homogenized with polypropylene pestle in Eppendorf plastic tubes (1.5 mL). Each sample was obtained in duplicate (8 to 10 larvae per sample). Total RNA was extracted with Reliaprep RNA tissue Miniprep System (Promega), according to the manufacturer’s instructions. Total RNA concentration was estimated using a Qubit™ 3.0 fluorometer (Invitrogen™) with Quantifluor RNA system (Promega). First-strand cDNAs were synthesized from each sample. The libraries were validated by real-time PCR using 2.5x reaction mixture for RT-PCR with EVA Green (Synthol, Russia) and primers for Illumina adaptors (Eurogen, Russia). Then they were sequenced on a HiSeq 2500 platform (Illumina, USA) using the HiSeq PE Rapid Cluster Kit v2 and HiSeq Rapid SBS Kit v2 (200 cycles, Illumina, USA) in the 100bp pair-end mode.

### Raw data processing, assembly, alignment and annotation

FastQC (version 0.11.8) software was used as quality control of raw sequence data (fastq files) to check all primary statistics (e.g. base sequence quality, GC content). Trimmomatic software (version 0.35) (Bolger et al. 2014) was used to cut adapter and other illumina-specific sequences from the reads (TruSeq3-PE), low average quality of reads within the window falls below a threshold in a sliding window approach (SLIDINGWINDOW:5:15) and not proper length of reads from raw data.

For genome assembly of *P. pembai*, we used MaSuRCA genome assembler (version 3.3.0) (Zimin et al. 2013) to create the set of genome scaffolds, of which only large ones (minimum 5000 bp length) remained as the final assembly. For assembler, we set 500 bp as mean and 50 bp as standard deviation of an insert size. Quast and BUSCO tools (versions 4.6.3 and 3.0.2, respectively) (Gurevich et al. 2013; Waterhouse et al. 2018) were used for quality assessment. According to Quast, we’ve got a set of 4662 scaffolds with total genome size of 122.9 Mb and N50 equal to 40.9 Kb of *P. pembai*. A tandem of RepeatModeler (version 1.0.11) and RepeatMasker (version 4.0.7) tools were used for identification of repeats and low-complexity regions, which were soft-masked for further gene prediction. This procedure was carried out with BRAKER tool (version 2.1.2) (Hoff et al. 2019) in *ab-initio* mode, that gave us a set of 15068 predicted genes. Functional annotation of corresponding protein models was performed using InterProScan pipeline (version 5.26-65.0) (Jones et al. 2014).

Bwa software with maximum exact match (mem) algorithm (Li and Durbin 2009) was used to map processed *P. vanderplanki* midge population data to the current version of *P. vanderlanki* reference genome assembly. Processed *P. pembai* midge population data was mapped to current *P. pembai* reference genome assembly based on Chikopa population. In order to estimate genetic distance between two species processed *P. pembai* midge population data was mapped to the current version of *P. vanderlanki* reference genome assembly.

Samtools (version 1.9) (Li et al. 2009) and Picard software (version 2.20.0) were used to convert sam format with deduplication step (picard) to sorted bam files. After collection insert size of pair end data, deduplication step was done by function MarkDuplicatesWithMateCigar with option minimum distance, which based on twice the 99.5% percentile of the fragment insert size.

Variant calling was performed with samtools mpileup (version 1.9) and bcftools call (or view) (version 1.9) options, where mpileup part generates genotype likelihoods at each genomic position with coverage and call part makes the actual calls.

Variant filtering was performed with GATK software (version 4.1.2.0), where positions or variants with depth (DP) in vcf files lower than 10 or mapping quality (QUAL) less than 20 are removed from vcf files. This step was done to get rid of the bias in the distribution of allele frequencies, as population data represent Pool-seq data.

### Phylogenetic analysis, polymorphisms estimation and evolution analysis

Phylogenetic analysis of different populations was done by MEGA7 software (Kumar et al. 2016) with maximum likelihood methods with bootstrap value (100) support. The consensus gene sequence was a sequence in which substitutions (SNP’s) with an allele frequency greater than 0.5 replaced the reference position.

Population diversity (π) was measured with custom script in R language. Interpopulation diversity was measured in the same manner as intrapopulation diversity, but in a pair-wide manner with different populations.

PoPoolation2 software tool (Kofler et al. 2011) was used to measure allele frequencies differences between populations and estimate population differentiation summary statistics (Fst) along the genome with sliding window approach. Input files for Popoolation2 software were synchronized files, which contain the allele frequencies after filtering for base quality for every population at every base in the reference genome. Synchronization was done by java-script implemented in PoPoolation2. Fst was calculated only for positions with coverage not lower than 9 reads.

Orthogroups for *P. vanderplanki* and *P. pembai* were obtained by OrthoFinder (version 2.3.11) (Emms and Kelly 2019). Sequence alignment was performed by mafft (v7.427) and subsequent filtering by trimal (version 1.4) software. To calculate dn/ds ratio codeml program from paml package (version 4.9) (Yang 2007) was used with the following parameters: runmode = −2, seqtype = 1, CodonFreq = 0, model = 0, NSsites = 0, fix_kappa = 0, kappa = 2, fix_omega = 0, omega = 0.5; for branch model we used the following options: runmode = 0, model = 2 and model = 0; and for site model: runmode = 0, model = 0, NSsites = 0 1 2 3 7 8, omega = 1.

### Analyses of differential expression at mRNA level

Transcriptomic data was represented by total RNA-seq derived from both species at three desiccation time points (D0, D24, D48). Reads mapped using hisat2 (version 2.1.0) (Kim et al. 2019) to genome assemblies of *P. vanderplanki* and *P. pembai* accordingly. Raw count data were then used as input into edgeR package (version 3.26.8) (Robinson et al. 2010) to analyse differentially expressed genes. The level of gene expression was compared at different hours of larvae desiccation.

## Author Contributions

D.P. and T.O. collected the midges, R.C. prepared samples, E.I.S., G.R.G, A.A.P. sequenced samples, O.S.K and A.V.C. assembled midges genomes, G.V.K., S.K.G., R.C., T.K., O.A.G and G.A.B. directed the research, N.M.S. analyzed the data and wrote the article. All authors contributed to the preparation of the article.

## Acknowledgments

This work was funded by Russian Scientific Foundation No. 20-44-07002. This work was also supported by the Institute of Agrobiological Sciences, National Agriculture and Food Research Organization (NARO) and in part by Grants-in-Aids from MEXT/JSPS KAKENHI (22128001, 23128512, 25128714), Japan. We are grateful to the University of Malawi for allowing us to use *P. pembai* specimens under MTA contract. We extend our gratitude to Prof. Augustine Eyiwunmi Falaye from the University of Ibadan, and IITA in Nigeria, for their kind cooperation. Finally, we thank Akihiko Fujita and Yoko Saito for their help in rearing the midges in the lab.

## References

1. Armitage PD, Pinder LC, Cranston PS. 2012. The Chironomidae: Biology and ecology of non-biting midges. Springer Science & Business Media

2. Boitard S, Kofler R, Françoise P, Robelin D, Schlötterer C, Futschik A. 2013. Pool-hmm: a Python program for estimating the allele frequency spectrum and detecting selective sweeps from next generation sequencing of pooled samples. Mol. Ecol. Resour. 13:337–340.

3. Bolger AM, Lohse M, Usadel B. 2014. Trimmomatic: a flexible trimmer for Illumina sequence data. Bioinformatics 30:2114–2120.

4. Chen L, DeVries AL, Cheng CH. 1997. Evolution of antifreeze glycoprotein gene from a trypsinogen gene in Antarctic notothenioid fish. Proc. Natl. Acad. Sci. U. S. A. 94:3811–3816.

5. Cornette R, Gusev O, Nakahara Y, Shimura S, Kikawada T, Okuda T. 2015. Chironomid midges (Diptera, Chironomidae) show extremely small genome sizes. Zoolog. Sci. 32:248–254.

6. Cornette R, Kikawada T. 2011. The induction of anhydrobiosis in the sleeping chironomid: current status of our knowledge. IUBMB Life 63:419–429.

7. Cornette R, Yamamoto N, Yamamoto M, Kobayashi T, Petrova NA, Gusev O, Shimura S, Kikawada T, Pemba D, Okuda T. 2017. A new anhydrobiotic midge from Malawi, *Polypedilum pembai* sp. n.(Diptera: Chironomidae), closely related to the desiccation tolerant midge, *Polypedilum vanderplanki* Hinton. Syst. Entomol. 42:814–825.

8. Desrosiers RR, Fanélus I. 2011. Damaged proteins bearing L-isoaspartyl residues and aging: a dynamic equilibrium between generation of isomerized forms and repair by PIMT. Curr. Aging Sci. 4:8–18.

9. Deviatiiarov R, Shagimardanova E, Kikawada T. 2017. Regulation of Gene Expression for L-Isoaspartyl O-Methyltransferases by Cis-Elements Associated with “Heat-Shock Polytene Chromosome Puffing Formation” in the Anhydrobiotic Midge. Bionanoscience 7:212–215.

10. Elliot MG, Crespi BJ. 2006. Placental invasiveness mediates the evolution of hybrid inviability in mammals. Am. Nat. 168:114–120.

11. Emms DM, Kelly S. 2019. OrthoFinder: phylogenetic orthology inference for comparative genomics. Genome Biol. 20:238.

12. Feakins SJ, Demenocal PB. 2010. Global and African regional climate during the Cenozoic. Cenozoic mammals of Africa :45–55.

13. Gurevich A, Saveliev V, Vyahhi N, Tesler G. 2013. QUAST: quality assessment tool for genome assemblies. Bioinformatics 29:1072–1075.

14. Gusev O, Cornette R, Kikawada T, Okuda T. 2011. Expression of heat shock protein-coding genes associated with anhydrobiosis in an African chironomid *Polypedilum vanderplanki*. Cell Stress Chaperones 16:81–90.

15. Gusev O, Suetsugu Y, Cornette R, Kawashima T, Logacheva MD, Kondrashov AS, Penin AA, Hatanaka R, Kikuta S, Shimura S, et al. 2014. Comparative genome sequencing reveals genomic signature of extreme desiccation tolerance in the anhydrobiotic midge. Nat. Commun. 5:4784.

16. Havird JC, Santos SR. 2014. Performance of single and concatenated sets of mitochondrial genes at inferring metazoan relationships relative to full mitogenome data. PLoS One 9:e84080.

17. Hinton HE. 1951. A new Chironomid from Africa, the larva of which can be dehydrated without injury. In: Proceedings of the Zoological Society of London. Vol. 121. Wiley Online Library. p. 371–380.

18. Hoff KJ, Lomsadze A, Borodovsky M, Stanke M. 2019. Whole-Genome Annotation with BRAKER. Methods in Molecular Biology [Internet]:65–95. Available from: http://dx.doi.org/10.1007/978-1-4939-9173-0_5

19. Innan H, Kondrashov F. 2010. The evolution of gene duplications: classifying and distinguishing between models. Nat. Rev. Genet. 11:97–108.

20. Jones P, Binns D, Chang H-Y, Fraser M, Li W, McAnulla C, McWilliam H, Maslen J, Mitchell A, Nuka G, et al. 2014. InterProScan 5: genome-scale protein function classification. Bioinformatics 30:1236–1240.

21. Keightley PD, Ness RW, Halligan DL, Haddrill PR. 2014. Estimation of the spontaneous mutation rate per nucleotide site in a Drosophila melanogaster full-sib family. Genetics 196:313–320.

22. Kelley JL, Peyton JT, Fiston-Lavier A-S, Teets NM, Yee M-C, Johnston JS, Bustamante CD, Lee RE, Denlinger DL. 2014. Compact genome of the Antarctic midge is likely an adaptation to an extreme environment. Nat. Commun. 5:4611.

23. Khare S, Linster CL, Clarke SG. 2011. The interplay between protein L-isoaspartyl methyltransferase activity and insulin-like signaling to extend lifespan in *Caenorhabditis elegans*. PLoS One 6:e20850.

24. Kikawada T, Nakahara Y, Kanamori Y, Iwata K-I, Watanabe M, McGee B, Tunnacliffe A, Okuda T. 2006. Dehydration-induced expression of LEA proteins in an anhydrobiotic chironomid. Biochem. Biophys. Res. Commun. 348:56–61.

25. Kim D, Paggi JM, Park C, Bennett C, Salzberg SL. 2019. Graph-based genome alignment and genotyping with HISAT2 and HISAT-genotype. Nat. Biotechnol. 37:907–915.

26. Kofler R, Pandey RV, Schlötterer C. 2011. PoPoolation2: identifying differentiation between populations using sequencing of pooled DNA samples (Pool-Seq). Bioinformatics 27:3435–3436.

27. Kondrashov FA. 2012. Gene duplication as a mechanism of genomic adaptation to a changing environment. Proc. Biol. Sci. 279:5048–5057.

28. Kumar S, Singh AK. 2017. Population Genetics of *Drosophila*: Genetic Variation and Differentiation among Indian Natural Populations of Drosophila ananassae. Zool. Stud. 56:e1.

29. Kumar S, Stecher G, Tamura K. 2016. MEGA7: Molecular Evolutionary Genetics Analysis Version 7.0 for Bigger Datasets. Mol. Biol. Evol. 33:1870–1874.

30. Li H, Durbin R. 2009. Fast and accurate short read alignment with Burrows–Wheeler transform. Bioinformatics 25:1754–1760.

31. Li H, Handsaker B, Wysoker A, Fennell T, Ruan J, Homer N, Marth G, Abecasis G, Durbin R, 1000 Genome Project Data Processing Subgroup. 2009. The Sequence Alignment/Map format and SAMtools. Bioinformatics 25:2078–2079.

32. McLachlan A. 1983. Habitat distribution and body size in rain-pool dwellers. Zool. J. Linn. Soc. 79:399–407.

33. Mendelson TC, Inouye BD, Rausher MD. 2004. Quantifying patterns in the evolution of reproductive isolation. Evolution [Internet] 58:1424. Available from: http://dx.doi.org/10.1554/03-632

34. Robinson MD, McCarthy DJ, Smyth GK. 2010. edgeR: a Bioconductor package for differential expression analysis of digital gene expression data. Bioinformatics 26:139–140.

35. Sogame Y, Kikawada T. 2017. Current findings on the molecular mechanisms underlying anhydrobiosis in *Polypedilum vanderplanki*. Curr Opin Insect Sci 19:16–21.

36. Tamura K, Battistuzzi FU, Billing-Ross P, Murillo O, Filipski A, Kumar S. 2012. Estimating divergence times in large molecular phylogenies. Proc. Natl. Acad. Sci. U. S. A. 109:19333–19338.

37. Watanabe M. 2006. Anhydrobiosis in invertebrates. Appl. Entomol. Zool. 41:15–31.

38. Watanabe M, Kikawada T, Minagawa N, Yukuhiro F, Okuda T. 2002. Mechanism allowing an insect to survive complete dehydration and extreme temperatures. J. Exp. Biol. 205:2799–2802.

39. Waterhouse RM, Seppey M, Simão FA, Manni M, Ioannidis P, Klioutchnikov G, Kriventseva EV, Zdobnov EM. 2018. BUSCO Applications from Quality Assessments to Gene Prediction and Phylogenomics. Mol. Biol. Evol. 35:543–548.

40. Yang Z. 2007. PAML 4: phylogenetic analysis by maximum likelihood. Mol. Biol. Evol. 24:1586–1591.

41. Zimin AV, Marçais G, Puiu D, Roberts M, Salzberg SL, Yorke JA. 2013. The MaSuRCA genome assembler. Bioinformatics 29:2669–2677.

