## Supplementary material for "Population genomics of two closely related anhydrobiotic midges reveals differences in adaptation to extreme desiccation": Supplementary material fig. S1-S4, table S1-4,7-8.docx

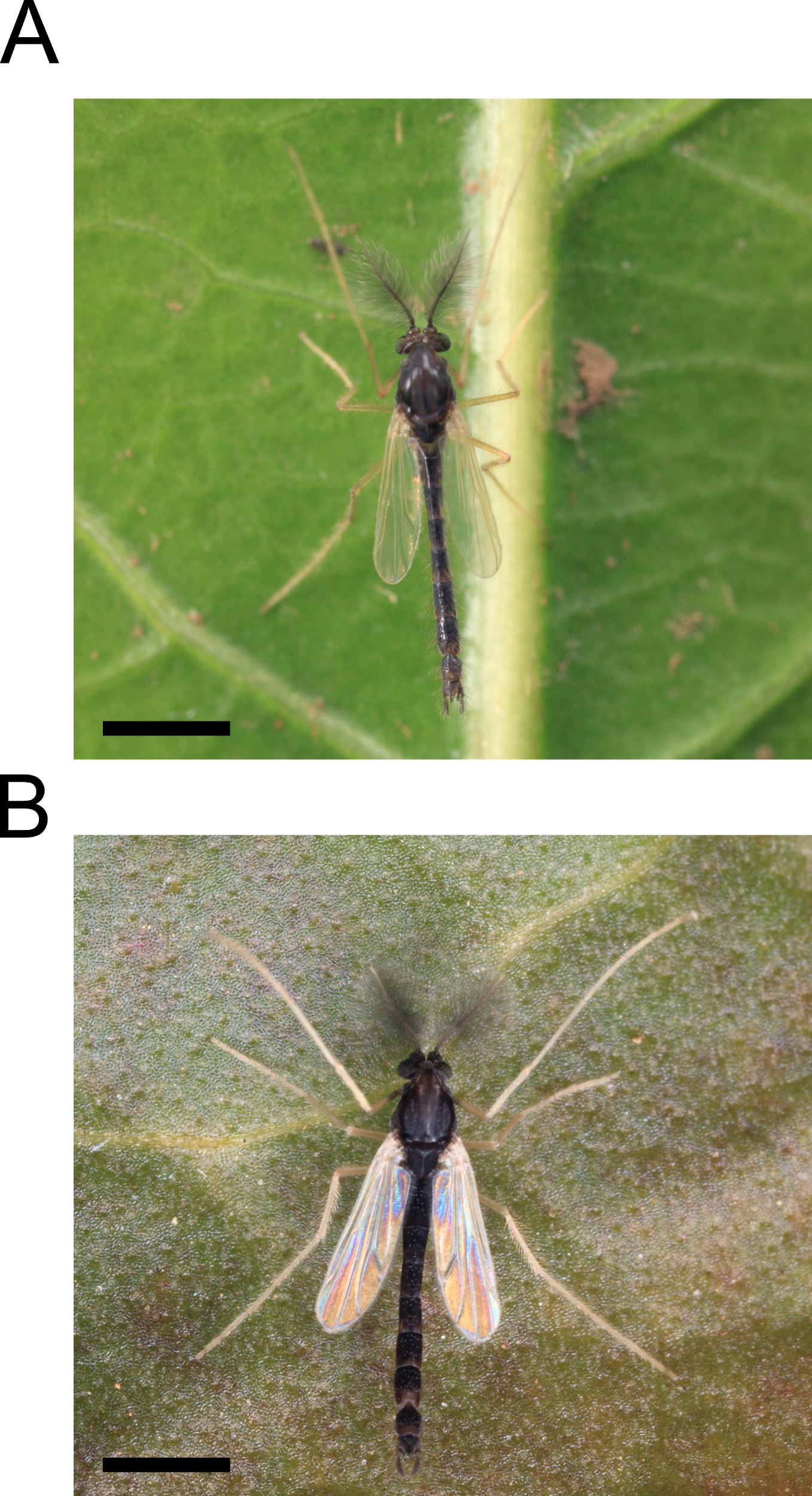

Fig. S1. Adult stages of *P. vanderplanki* (A) and *P. pembai* (B). Scale bars represent 1 mm.

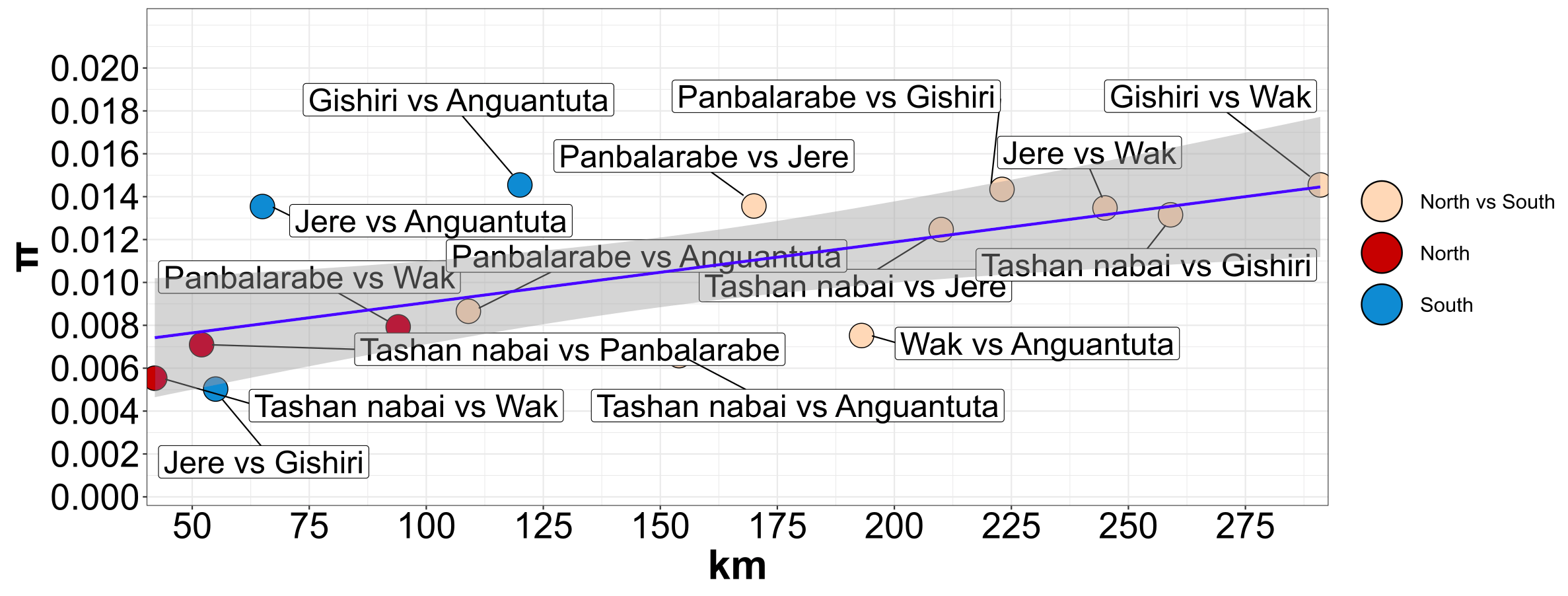

Fig. S2. Pairwise nucleotide diversity between all *P. vanderplanki* studied populations as a function of geographic distance (km).

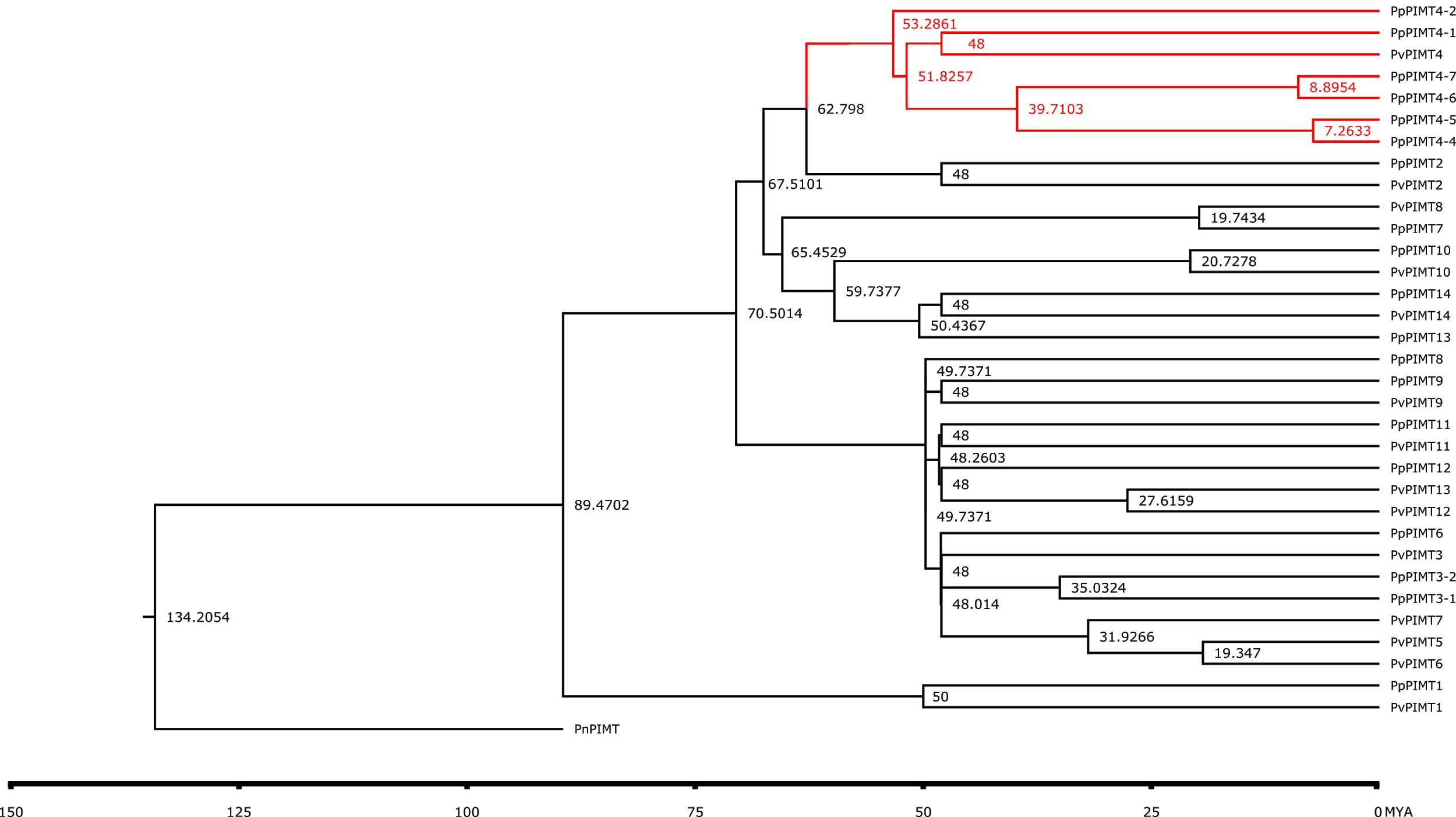

Fig. S3. Time divergence of PIMT paralogs using Reltime-ML method.

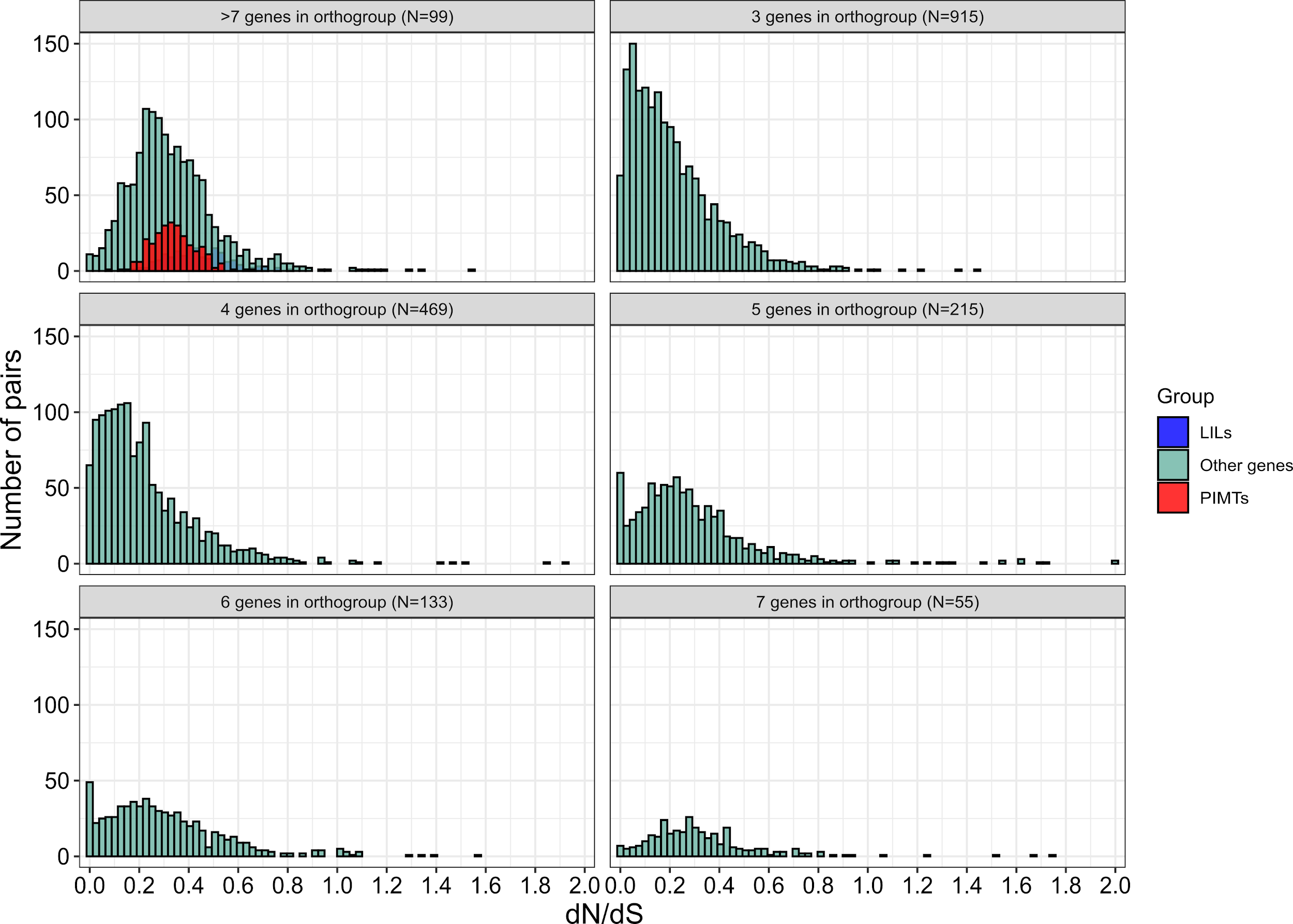

Fig. S4. Distribution of dn/ds in different orthologous groups size.

Table S1. Pool-seq data analysis summary

| Population | Total number of reads | Number of reads after trimming | Mapping  rate, % | Average coverage |
| --- | --- | --- | --- | --- |
| Tashan nabai | 6806884 | 6456690 | 98.87 | 12.03 |
| Panbalarabe | 8233398 | 7841488 | 98.91 | 14.29 |
| Wak | 9644900 | 9295736 | 98.76 | 17.16 |
| Jere | 6029944 | 5734942 | 98.01 | 10.23 |
| Gishiri | 8621858 | 8218478 | 98.81 | 14.98 |
| Anguantuta | 21303324 | 17243054 | 81.43 | 10.34 |
| Сhikopa | 114356234 | 100472154 | 90.1 | 163.5 |

Table S2. The statistics of the assembled genomes of *P. vanderplanki* and *P. pembai.*

|  | *P. pembai* | *P. vanderplanki* |
| --- | --- | --- |
| Number of scaffolds | 4662 | 4 |
| Genome size, Mbp | 122 | 120.4 |
| Coverage | ~110X | ~450X |
| N50 (scaffold) | 40.9 kbp | 34 Mbp |
| The amount of N per 100 kbp | 208.78 | 1699.08 |
| BUSCO* | C: 95.1%;  D:1.5%;  F: 2.5%;  M: 2.4% | C: 96.8%;  D: 0.9%;  F: 1.7%;  M: 1.5% |
| Genomic G+C content, % | 28.6 | 28.1 |
| Predicted numbers of protein-coding genes | 15068 | 17852 |
| Percentage of genes in orthogroups | 93.8 | 91.9 |
| Genes common for both species (number of 1:1 orthologues) | 10625 | 10625 |

Note. —Letters in BUSCO correspond to complete, duplicated, fragmented and missing.

Table S3. Total RNA-seq summary characteristics for *P. pembai*

| Sample | Total number of reads (2x100) | Overall alignment rate, % | Uniquely mapped reads, % | Multi mapped reads, % |
| --- | --- | --- | --- | --- |
| D0 (first replica) | 133996878 | 19.31 | 11.13 | 3.63 |
| D0 (second replica) | 124822970 | 17.33 | 10.19 | 2.89 |
| D24 (first replica) | 128101442 | 16.75 | 9.37 | 2.94 |
| D24 (second replica) | 146232426 | 16.00 | 8.98 | 2.79 |
| D48 (first replica) | 129063180 | 18.10 | 10.01 | 03.05 |
| D48 (second replica) | 159699192 | 16.65 | 9.50 | 2.90 |

Table S4. Intrapopulation and interpopulation genetic distances of Nigerian populations. Red, populations from northern Nigeria; blue, populations from southern Nigeria.

| π | Tashan nabai | Panbalarabe | Wak | Jere | Gishiri | Anguantuta |
| --- | --- | --- | --- | --- | --- | --- |
| Tashan nabai | 0.004 |  |  |  |  |  |
| Panbalarabe | 0.007 | 0.007 |  |  |  |  |
| Wak | 0.005 | 0.008 | 0.005 |  |  |  |
| Jere | 0.012 | 0.013 | 0.013 | 0.004 |  |  |
| Gishiri | 0.013 | 0.014 | 0.014 | 0.005 | 0.004 |  |
| Anguantuta | 0.009 | 0.011 | 0.01 | 0.014 | 0.015 | 0.0057 |

Table. S7. πN/πS ratio of PvPIMT genes in all Nigerian midge populations (except Anguantuta).

Asterisks in table 13 mean that πN or πS of this particular PIMT is zero, so the πN/πS of this PIMT

cannot be measured.

|  | πN/πS | | | | |
| --- | --- | --- | --- | --- | --- |
| Genes | Tashan nabai | Panbalarabe | Jere | Gishiri | Wak |
| PvPIMT1 | * | 0,05 | * | * | * |
| PvPIMT2 | 0,068 | 0,04 | 0,594 | 0,044 | 0,051 |
| PvPIMT3 | * | 0,019 | * | * | * |
| PvPIMT4 | 0,046 | 0,206 | * | 0,03 | 0,065 |
| PvPIMT5 | 0,035 | 0,087 | 0,06 | 0,049 | 0,074 |
| PvPIMT6 | 0,113 | 0,11 | 0,033 | 0,056 | 0,098 |
| PvPIMT7 | 0,02 | 0,032 | * | * | 0,129 |
| PvPIMT8 | * | 0,372 | 0,06 | 0,395 | 0,545 |
| PvPIMT9 | 0,078 | 0,021 | * | 0,082 | 0,065 |
| PvPIMT10 | * | * | * | 0,02 | * |
| PvPIMT11 | 0,105 | 0,041 | * | 0,024 | 0,037 |
| PvPIMT12 | 0,035 | 0,066 | 0,02 | 0,043 | 0,045 |
| PvPIMT13 | 0,018 | 0,014 | * | 0,043 | 0,037 |
| PvPIMT14 | 0,096 | 0,084 | 0,029 | 0,059 | 0,051 |

Table S8. Genetic distance between *P. vanderplanki* and *P. pembai.* Chikopa is the name of *P. pembai* population

| Populations | π (all genome) | π (chr4) | π (chr3) | π (chr2) | π (chr1) | Mean Fst along full genome |
| --- | --- | --- | --- | --- | --- | --- |
| Tash vs. Chikopa | 0.078 | 0.098 | 0.079 | 0.076 | 0.076 | 0.9 |
| Pan vs. Chikopa | 0.076 | 0.1 | 0.077 | 0.074 | 0.074 | 0.85 |
| Jere vs. Chikopa | 0.08 | 0.1 | 0.081 | 0.078 | 0.078 | 0.91 |
| Gishiri vs. Chikopa | 0.079 | 0.098 | 0.079 | 0.077 | 0.075 | 0.9 |
| Wak vs. Chikopa | 0.077 | 0.096 | 0.077 | 0.075 | 0.076 | 0.88 |
| Anguantuta vs. Chikopa | 0.079 | 0.098 | 0.079 | 0.077 | 0.076 | 0.89 |
